# From Pigments to Precision: Exploring Genetic Transformation and Genome Editing in Wheat and Barley

**DOI:** 10.1101/2024.12.03.626565

**Authors:** Manas Ranjan Prusty, Arava Shatil-Cohen, Rakesh Kumar, Davinder Sharma, Anna Minz-Dub, Smadar Ezrati, Avigail Hihinashvili, Amir Sharon

## Abstract

Genetic engineering of wheat is complex due to its large genome size, the presence of numerous genes with high sequence similarities, and a multitude of repetitive elements. In addition, genetic transformation of wheat has been difficult, mainly due to poor regeneration in tissue cultures. Recent advances in plant biotechnology, particularly the use of the regenerative genes GROWTH-REGULATING FACTOR (*GRF*) and GRF-INTERACTING FACTOR (*GIF*), have provided new tools for wheat transformation and regeneration. Another transformative tool is the RUBY system, that involves genetic engineering of three betalain biosynthesis genes, providing a noninvasive, visually detectable red pigment. In this study, we used the *GRF4-GIF1* chimera along with the RUBY system to advance transformation and gene editing in wheat and barley. The GRF4-GIF1 chimera significantly aided wheat regeneration; however, it had an opposite effect in barley, where it inhibited the regeneration process. Therefore, we primarily generated RUBY transgenic barley lines using constructs that did not include the GRF4-GIF1 chimera. Additionally, we used the RUBY cassette for fast assessment of gene editing by knockingout the first betalain biosynthetic gene in RUBY-positive transgenic wheat plants, resulting in a change of leaf color from red to green. The edited RUBY wheat lines lost more than just the red color. They also lost betalain-related traits, such as being less likely to get leaf rust (*Puccinia triticina*) and salt stress. Importantly, the loss of RUBY did not affect plant viability, making it a useful tool for genome editing and a viable alternative to destructive methods.

## Introduction

Wheat has been recalcitrant to genetic transformation relative to the two other staple crops, maize and rice (Hayta *et al*., 2021). Two main issues that have limited wheat transformation efficiency are the large and complex genome and the difficulties in regenerating whole plants from transformed tissues (Debernardi *et al*., 2020). Various methods have been employed for wheat transformation, including agrobacterium-mediated transformation, particle bombardment (biolistics), and in planta methods (Yu *et al*., 2024). Particle bombardment is widely used across different wheat varieties; however it often results in multiple gene copies being inserted (Tassy *et al*., 2014; Tanaka *et al*., 2022; Liang *et al*., 2018), which complicates the analysis of transgenic lines. In planta methods, although promising, are still under development for wheat and have not yet been widely adopted (Hamada *et al*., 2017). The ability of Agrobacterium-mediated transformation to stably introduce a single or low number of copies is particularly valuable, but its efficiency varies significantly depending on the wheat cultivar (Zhang *et al*., 2019). Regenerative genes have been tested as a means to overcome the low rates of wheat tissue regeneration. Researchers have shown that expressing a fusion protein that combines wheat GROWTH-REGULATING FACTOR 4 (GRF4) with its co-factor GRF-INTERACTING FACTOR 1 (GIF1) (Debernardi et al., 2020; Biswal et al., 2023) and the PANICLE1 (*TaLAX1*) gene (Yu et al., 2024) enhances regeneration in genotypes of wheat that are otherwise difficult to transform. Similarly, overexpression of the wheat TaWOX5 gene has significantly improved transformation efficiency across wheat varieties and other cereal plant species (Wang et al., 2022). The RUBY gene system has emerged as a useful reporter in plant research (He et al., 2020). The “2A” peptide helps this system use the three betalain biosynthetic genes—P450 oxygenase (*CYP76AD1*), l-DOPA 4,5-dioxygenase (*DODA*), and glucosyl transferase (*GT*)—to make betalain, a beat red pigment. It has been used to advance transformation of plants like rice, Arabidopsis, and cotton and proven useful for other purposes, such in visualization of protein-protein interactions (Pierroz, 2023) and accurate identification of haploids in maize and tomato (Wang *et al*., 2023). Furthermore, RUBY played a critical role in gene stacking and was utilized as a split selectable marker system to boost co-transformation efficiency in model plants like Arabidopsis and poplar (Chen *et al*., 2023; Pierroz, 2023).

Here, we expanded the application of RUBY for genome manipulation and used it for advancing transformation and gene editing in wheat and barley **(Figure 1, Figure S1)**. First, we used RUBY for improvement of transformation efficiency in wheat cv. Fielder, the most commonly used cultivar for wheat transformation, cv. Chinese spring, the first fully sequenced wheat cultivar, and cv. Gadish and cv.Bobwhite a widely used commercial wheat cultivar, as well as in barley. We then used RUBY to evaluate and adjust CRISPR/Cas9 editing in wheat, showcasing the effective editing of a transgenic RUBY cassette as a control in gene editing experiments

**Figure 1.**
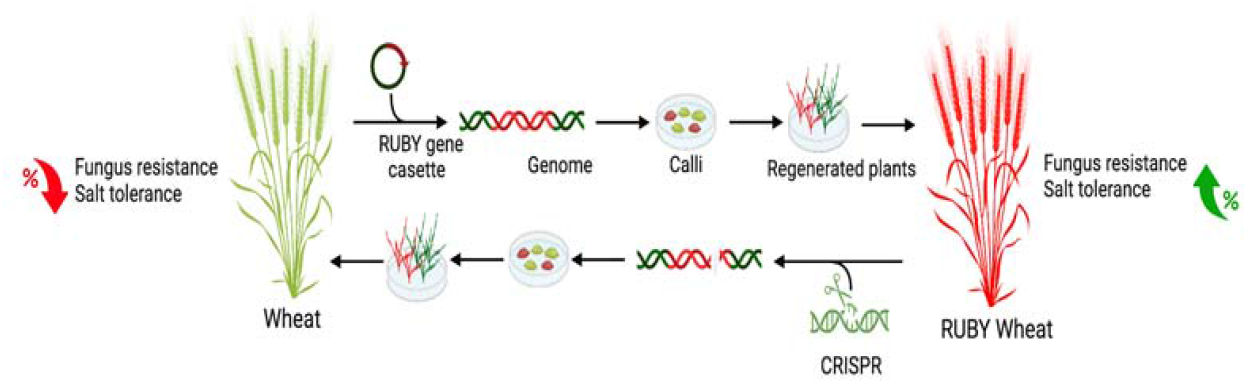
Schematic outline of the use of RUBY in wheat transformation and gene editing. Expression of the RUBY cassette in wheat cv. Fielder results in transgenic plants with a distinct red pigment. Successful editing of the RUBY plants results in green plants. Along with the loss of the red pigment, the edited lines also lose the betalain-associated traits, including reduced sensitivity to leaf rust and salt stress.

The loss of RUBY pigment in edited plants was accompanied by loss of betalain-associated traits, including reduced susceptibility to leaf rust (*Puccinia triticina Eriks*., *Pt*) and salt stress, demonstrating that the editing process affected both the visible phenotype as well as the RUBY-related traits. Our study demonstrates the potential of RUBY as a useful tool for calibrating and advancing transformation and CRISPR/Cas9 editing protocols in plants.

## Results

### Selection of promoters

To select suitable promoters for gene expression in wheat, we tested the activity of five commonly used promoters: maize ubiquitin (*ZmUbi*), rice actin (*OsActin*), cauliflower mosaic virus 35S (*CaMV35S*), wheat actin (*TaActin*), and wheat ubiquitin (*TaUbi*). We linked each promoter to the wheat codon-optimized enhanced green fluorescent protein (*eGFP*) gene **(Figure S2a)** and cloned the constructs into the pJET1.2 vector (Thermo Fisher Scientific). We transformed the resulting plasmids into wheat protoplasts via PEG-mediated transformation, and into wheat immature embryos using biolistic delivery with gold particles. We evaluated the relative strength of the different promoters by monitoring GFP fluorescence levels **(Figure S2b)**. Initially, we tested the construct with the *CaMV35S* promoter, which has been widely used in various plant species including wheat. Consistent with Rather et al. (2022), we found that a linearized plasmid enhanced protoplast transformation efficiency compared to circular DNA; however when we used biolistic transformation, the GFP signal was observed only in embryo that were transformed with a circular plasmid **(Figure S2c)**. To change the rest of the plasmids, we used linearized plasmids to change protoplasts through PEG and circular plasmids to change embryos completely.

Among the five promoters, the *ZmUbi, CaMV35S*, and *OsActin* promoters produced consistent and robust GFP fluorescence in both protoplasts and wheat embryos **(Figure 2**,**Figure S2b)**. However, we could not detect a GFP signal in protoplasts or embryos that were transformed with either of the two wheat promoters, *TaActin* and *TaUbi*, indicating unexpected low transcriptional activity of these wheat endogenous promoters (Figure 2). Based on these findings, we chose the *ZmUbi, CaMV35S*, and *OsActin* promoters for construction of our transformation and editing plasmids.

**Figure 2.**
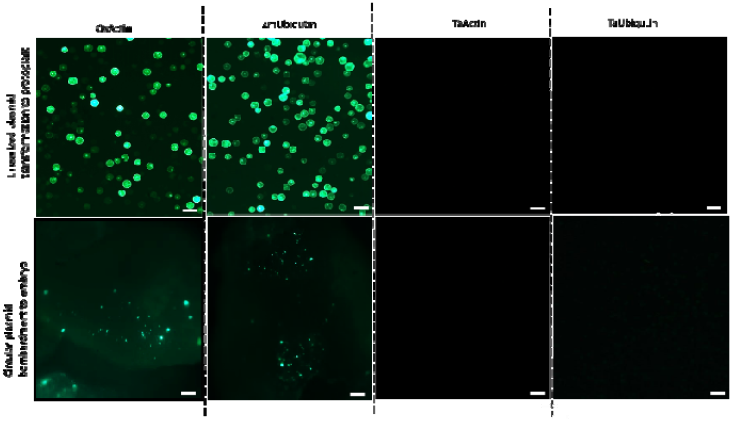
Screening of promoters by expression in wheat protoplasts and embryos. Protoplasts were transformed with linearized DNA by PEG-mediated transformation (top images); embryos were transformed with circular DNA by biolistic transformation (bottom images). Promoters activity was evaluated by monitoring GFP fluorescence intensity using a fluorescent microscope. Scale bar in upper and lower images = 100 and 500 µm, respectively.

### Generation of wheat and barley transgenic RUBY lines

To generate betalain producing plants, we cloned a cassette containing the Beta vulgaris betalin biosynthesis genes *CYP76AD1, DODA*, and *GT* under the control of the *CaMV35S* promoter into the ICCR2 plasmid (Sharma et al., 2024), which includes the GRF4-GIF1 chimera (Debernardi *et al*., 2020) driven by the *ZmUbi* promoter, along with the *BAR* gene that confers resistance to the herbicide Basta **(Figure 3a)**. We transformed immature embryos of wheat Fielder with the resulting (ICCR2:RUBY) and the ICCR2 (control) plasmids **(Figure 3a,b)**, obtained independent transgenic lines and confirmed the presence and expression of the *BAR* and *GRF4-GIF1* genes in these lines by RT-PCR **(Figure 3c-d, Figure S1)**. The ICCR2:RUBY plasmid successfully transformed and expressed the RUBY cassette in approximately 47% of the regenerated plants**(Figure S3e)**. All parts of the transgenic RUBY lines were red, including the spikelet and roots **(Figure 3d)**. We used digital droplet PCR (ddPCR) to evaluate the transgene copy number in four transgenic lines each of plants expressing the ICCR2 or ICCR2:RUBY. The copy number ranged from 2 (homozygote single insertion) to 6 in the control transgenic lines and from 2 to 12 in Ruby transgenic lines **(Figure S4)**.

**Figure 3.**
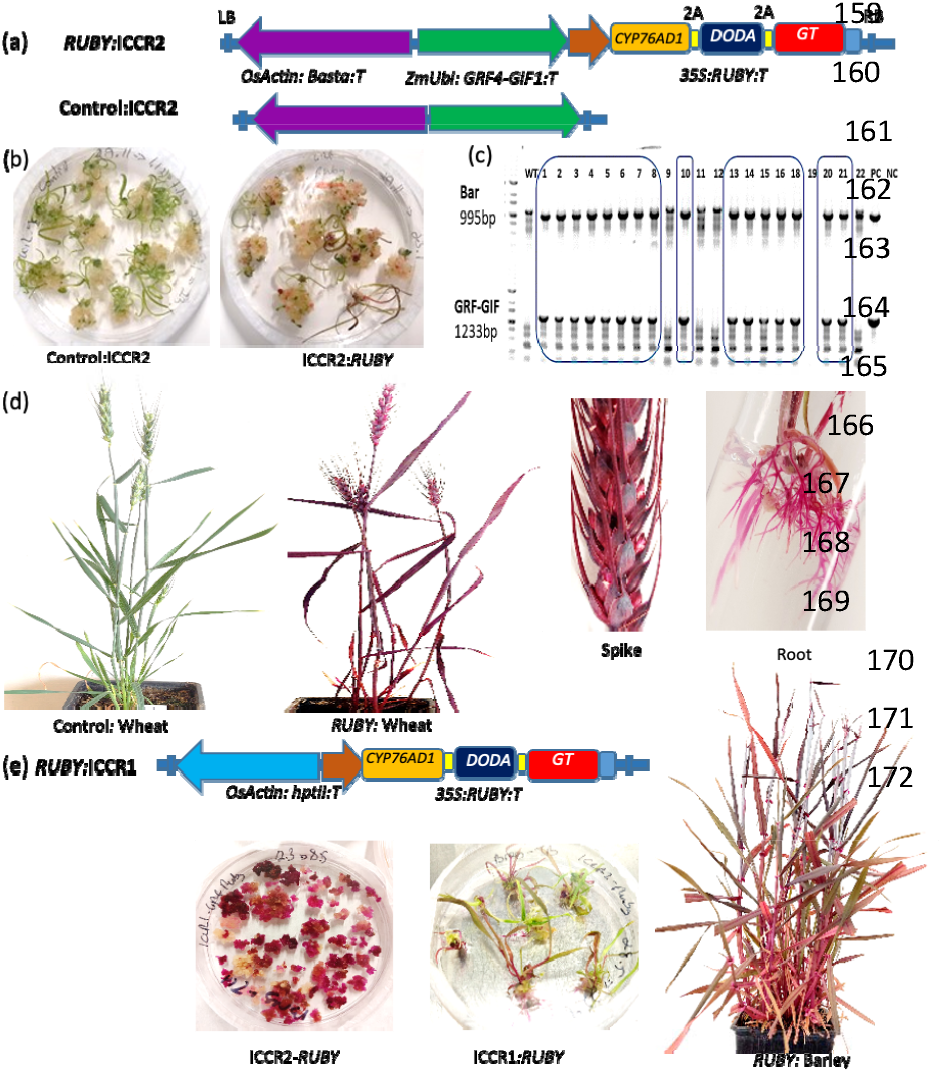
Development of RUBY wheat and barley transgenic lines. (a) Illustrates the schemes of the ICCR2:RUBY and ICCR2 plasmids. Both plasmids include the BAR gene under the *OsAct* promoter (purple) and the *GRF4-GIF1* chimera under the *ZmUbi* promoter (green). The RUBY cassette includes the *CYP76AD1, DODA*, and *GT* genes, separated by the 2A peptide, all under the CaM35S promoter. LB and RB denote the left and right borders of the T-DNA region, respectively. (b) Transforming wheat calli with ICCR2 (left) or ICCR2:RUBY (right) results in regeneration (c) RT-PCR analysis of transgenic T0 plants using primers for amplification of the *BAR* (995 bp) and *GRF4-GIF1* (1233 bp) genes; WT; wild type, PC; positive control (ICCR2 plasmid), NC; negative control (water). (d) Transgenic wheat lines transformed with ICCR2 (left) and ICCR2:RUBY (right). RUBY is expressed in all parts of the plants, including spikelets and roots. (e) Barley transformation. Transformation with the ICCR2:RUBY plasmid resulted in red callus but low regeneration (left), transformation with the ICCR1:RUBY plasmid that does not contain the *GRF4-GIF1* chimera resulted in high regeneration rates and transgenic plants.

We extended the use of the RUBY reporter to calibrate transformation in three additional wheat varieties: Bob White, Chinese Spring (CS), and Gadish, a widely used commercial cultivar **(Figure S3, a-d)**. The transformation efficiency varied among these varieties, with success rates of 20% for Bob White, 8% for CS, and 7.5% for Gadish **(Figure S3, e)**. The fact that this works shows how useful the GRF4-GIF1 chimera and the RUBY cassette are for quickly testing and developing transformations in a variety of wheat genotypes.

Unexpectedly, while significantly enhancing regeneration of wheat calli, the *GRF4-GIF1* chimera had a negative effect on regeneration in barley. The initial calli turned red, indicating successful transformation; however, the majority of the calli remained undifferentiated, with less than 10% final regeneration rates **(Figure 3e, Figure S3e)**. In contrast, when we used the ICCR1 plasmid, which does not include the *GRF4-GIF1* chimera, regeneration efficiency reached 85%, and approximately 53% of the regenerated plants exhibited the RUBY phenotype **(Figure 3e, Figure S3e)**. Hence, using the RUBY cassette, we managed to quickly identify the negative effect of the *GRF4-GIF1* chimera on barley transformation and pin point it to the stage of callus regeneration.

### Knocking out the RUBY cassette

To knock out the RUBY cassette, we designed a guide RNA (gRNA) that targets CYP76AD1, the first gene in the RUBY cassette **(Figure 4a)**, and cloned it into the JD633 binary plasmid (Addgene 160392). We chose the T1 RUBY transgenic line-2 (RUBY-2) **(Figure S1)** for the transformation experiments because it has a single, homozygote insertion of the transgene **(Figure S4)**. The T0 plants that were grown from the edited RUBY-2 line had a chimeric phenotype, which means that some of their tissues were colored both green and red **(Figure 4b, Figure S5)**. In the T1 generation, the plants segregated into entirely green or entirely red plants **(Figure S6)**. To check the editing events, we chose eight green T1 plants and three red T1 plants, each from a separate transformation event. We then amplified and sequenced a 500-bp region that covered the gRNA target site in the CYP76AD1 gene. As a control, we sequenced DODA, the second gene in the RUBY cases. Our research showed that four different editing events happened near the PAM site of the CYP76AD1 gene in plants that were green (Figure 4e, Figure S6). To verify the editing events, we selected eight green and three red T1 plants, each from an independent transformation event, and amplified and sequenced a 500 bp region spanning the gRNA target site within the *CYP76AD1* gene. As a control, we sequenced *DODA*, the second gene in the RUBY cassette **(Figure S7)**. We identified four distinct editing events (designated E1-E4) near the PAM site of the *CYP76AD1* gene in plants exhibiting the green phenotype **(Figure 4e, FigureS6)**. In E1, a single base pair insertion caused a frameshift mutation, leading to the production of a truncated protein with 205 amino acids, compared to the 1,316 amino acids CYP76AD1 protein. The E2 mutation caused a 3-base pair deletion, which lost one amino acid and made the protein slightly shorter, with 1,315 amino acids instead of 1,317. This showed that amino acid 1316 is necessary for the enzyme activity of CYP76AD1. E3 and E4 featured 8 and 12 base pair deletions that yielded truncated proteins of 203 and 493 amino acids, respectively **(Figure S8)**. The overall editing efficiency was approximately 33.3% **(Figure S6e)**. There were no changes in the control *DODA* gene in any of the plants nor in the *CYP76AD1* gene in plants with a red phenotype **(Figure S7)**.

**Figure 4.**
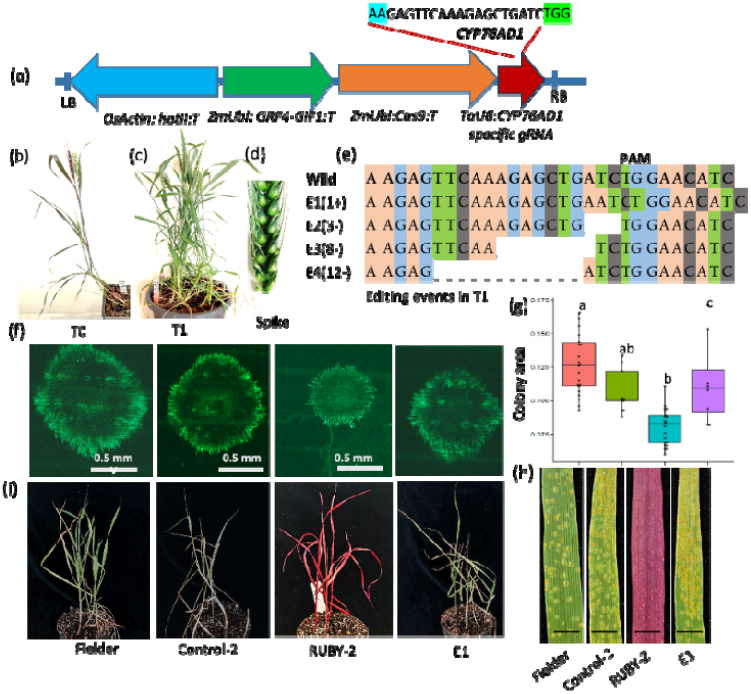
Editing of the *CYP76AD1* gene and its effects on betalain production and plants sensitivity to leaf rust and salt stress. (A) Diagram of the gene editing vector JD633, which includes the gRNA sequence targeting the *CYP76AD1* gene. The start site of gRNA is marked in blue, the PAM site in green. (b) A T0 edited line showing a chimeric phenotype characterized by both red and green coloration. See also Figure S4. (c) Whole plant and a spikelet of a green, RUBY edited T1 plant. (e) Summary of editing events: “+” denotes insertions, “–” indicates deletion. (f) Loss of betalain production correlates with reduced resistance in edited lines, as shown by fluorescence images of infection sites at 7 days post-inoculation (dpi), with fungal hyphae stained using WGA-FITC. (g) Box plot and ANOVA results with Tukey’s pairwise comparison for the pustule colony area. The different colors used are red for Fielder, green for control-2, blue for RUBY-2, and purple for E1. Different letters indicate statistically significant differences (P ≤ 0.01). (h) Infected leaves 7 dpi. Scale bar in all images =1mm. (i) Sensitivity to salt stress. Plants were evaluated after irrigation with 200 mM NaCl for 14 days. See also Figure S9.

Betalaines have been reported to reduce the susceptibility of transgenic RUBY tobacco plants to *Botrytis cinerea* (Polturak *et al*., 2017) and contribute to salt stress tolerance in *Disphyma austral* (Jain and Gould, 2015). So, we looked at how vulnerable our RUBY and RUBY edited wheat lines were to salt stress and the leaf rust pathogen *Puccinia triticina* (Pt). We included a transgenic line that was changed with the GRF4-GIF1 cassettes without RUBY because the RUBY transgenic lines have the GRF4-GIF1 chimera. We also included the native Fielder cultivar.

We evaluated the fungal infection by visual assessment of pustule size and leaf coverage, and by staining leaves with WGA-FITC and examining them under a fluorescent microscope to measure the area occupied by the fungus under infection sites. The RUBY plants exhibited slower disease progression, with a smaller fungal occupied area and fewer pustules compared with the control, whereas the RUBY edited line was as sensitive as the control plants **(Figure 4f-h)**. For salt stress, we visually assessed the plant phenotype after 14 days of irrigation with 200 mM NaCl. The RUBY edited line was just as sensitive to salt stress as the control plants. Older leaves turned a bright yellow color and young leaves rolled up, but the leaves on the unedited RUBY transgenic lines were barely deformed **(Figure 4i, Figure S9)**. Thus, just as the RUBY-edited lines lost their pigmentation, they also lost their enhanced resistance to fungal infection and salt stress.

## Discussion

Genetic transformation in plants relies on various types of markers to identify and select successfully transformed individuals. Among the most commonly used markers are antibiotics and herbicide resistance genes, visual markers, and molecular markers, each with specific advantages and limitations (Miki and McHugh, 2004). The advantage of visible phenotype markers is their ability to track the transformation process at all stages. Fluorescent proteins, such as GFP, are popular visible markers and have been successfully applied for assessment of transgenic lines in a wide range of plant species (Chiu *et al*., 1996; Sparkes *et al*., 2006; Jung *et al*., 2011; Omar *et al*., 2019; Jin *et al*., 2012; Dunbar *et al*., 2022). However, unlike many other plant species, achieving stable GFP expression in transgenic wheat and barley has proven challenging. While GFP signals are readily detectable in transient assays such as protoplast transformation or in calli, generating a transgenic wheat plant that expresses GFP remains difficult (Hamada *et al*., 2017). Indeed, we failed to generate transgenic wheat lines that express GFP; while we observed a strong signal in calli, we could not detect the GFP signal in regenerated plants, despite the presence of the *eGFP* gene **(Figure S10)**. In this context, the RUBY cassette represents a promising and attractive alternative; the color is readily detected by the bare eye already at the early stages of the tissue culture, it can be followed during all stages of plant regeneration, and is well expressed in the mature wheat and barley transgenic lines. Unlike antibiotic or herbicide resistance, RUBY provides a safe and environmentally friendly option, meeting the increasing demand for sustainable transformation techniques. Additionally, this marker system reduces the need for extensive PCR testing and sequencing.

The *PDS* (phytoene desaturase) gene is commonly used to assess gene silencing and gene editing efficacy, as disruption of PDS leads to a distinct albino phenotype (Banakar *et al*., 2020; Siddappa *et al*., 2023) However, blocking of chlorophyll biosynthesis can adversely affect plant growth and survival, which restricts the use of *PDS* to local effects. RUBY offers a viable alternative; our RUBY edited plants did not show any developmental change, the plants looked healthy throughout their life cycle and produced normal amounts of seeds **(Figure S6)**, and editing of RUBY reverted the phenotype to the wild type. Additionally, we did not observe any negative effect of RUBY on tissue regeneration and transformation rates. The RUBY system not only helped in quick validation of the edited plants, it also provided a framework for transgene editing strategies, especially when copy number is crucial. Furthermore, in polyploid plant species, such as wheat, editing endogenous versus transgenes requires different approaches. For endogenous genes with multiple copies, the design of the gRNA must take in consideration specificity and off-target effects, and for loss of function, all copies must be edited. For a heterologous transgene the off-target issues are less critical since it is foreign to the wheat genome, making copy number estimation more important for successful editing. Therefore, in our study, we specifically selected a RUBY transgenic line that contains a single homozygote RUBY line.

Previous studies have demonstrated that betalains can lessen the susceptibility of tobacco plants to the necrotrophic fungus Botrytis cinerea (Polturak *et al*., 2017). Here, we showed that the RUBY lines reduced wheat susceptibility to *Pt*, a biotrophic pathogen that causes leaf rust of wheat. We also found that the betalain producing plants were less sensitive to salt stress, an important phenomenon that has been previously recognized in *Disphyma austral* (Jain and Gould, 2015). Along with the loss of color, the RUBY-edited lines also lost the leaf rust and salt tolerance, confirming the role of betalains in these stress responses. Further research could explore the broader stress resistance capabilities of RUBY lines, potentially testing them against a range of abiotic and biotic stresses.

We made two additional key observations during this study. First, we noticed that protoplast transformation was more efficient with linearized DNA, whereas circular DNA was required for biolistic transformation of embryos. This difference likely arises from the distinct methods of DNA uptake: protoplasts, lacking cell walls, more easily integrate linearized plasmids, leading to higher gene expression, whereas biolistic delivery into intact cells possibly favors supercoiled plasmids due to their higher stability. The second difference was the effect of the *GRF4-GIF1* chimera on regeneration in wheat and barley. While the *GRF4-GIF1* chimera greatly improved regeneration in wheat, it inhibited tissue regeneration in barley. The red color of barley calli indicated that transformation was highly efficient; however the cells failed to regenerate, possibly due to impairment of the native regenerative pathways by the wheat *GRF4-GIF1* genes.

Our study underscored the RUBY system’s potential as a valuable tool for genetic transformation and gene editing, albeit with certain limitations that require attention. One limitation pertains to the efficient use of RUBY-expressing protoplasts for detecting gene editing events. While protoplast systems are well-established for assessing gRNA editing efficiency (Yang *et al*., 2024; Lin *et al*., 2018) and guide RNAs have been used to target GFP in protoplasts (Najafi *et al*., 2023; Zhang *et al*., 2019). RUBY color is highly sensitive to pH changes (Polturak *et al*., 2017). During protoplast isolation, chemicals and conditions often alter pH levels, causing the RUBY pigment to degrade or change color, resulting in some protoplasts appearing green instead of the expected red color **(Figure S11)**. Another limitation is the time required for the development of the RUBY color. The pigment typically becomes visible in calli about three weeks after transformation, which is significantly slower than GFP expression in transient assays. Overall, despite these limitations, RUBY proves to be a versatile, visually detectable marker for plant transformation and gene editing in wheat and barley. RUBY could be used as a reporter gene in more crop plants in the future. It could also be used to make plants more resistant to biotic and abiotic stresses, which would make it even more useful in plant biotechnology.

### Experimental procedure

#### Protoplast Transformation

Protoplasts were isolated from 1-week-old seedlings of wheat cv. Fielder following (Brandt *et al*., 2020) with some modification. We rinsed approximately 200 mg of young leaves with distilled water, chopped them into 0.5-mm strips, and incubated them in a digesting enzyme solution at 27°C in the dark for 2-3 hours. We diluted the mixture with an equal volume of W5 solution (MES 2mM, NaCl 154mM, CaCl2 125mM, KCl 5mM). We then filtered the mixture through a 100-m cell strainer and centrifuged it at 100g for 2 minutes to extract the protoplasts. After removing the supernatant, we washed the pellet with W5A solution (Glucose 5mM, MES 2mM, NaCl 154mM, CaCl2 125mM, KCl 5mM), and layered it onto a 21% sucrose solution. s centrifuged at 720 g for 15 minutes, the viable protoplasts were carefully collected, and their density was adjusted to 0.7–1.0 × 10^6 cells/ml in MMG solution (D mannitol 0.5M, KCl 20mM, MES 4mM). For GFP plasmid transformation, 15 µg of the plasmid were mixed with 200 µl of protoplast solution and 230 µl of PEG solution (40% PEG 4000, 0.2M D-mannitol, 0.1M CaCl2), and the mixture was incubated at room temperature for 30 minutes. We stopped the reaction by adding 1 ml of W5 solution, resuspended the protoplasts in 500 ul of fresh W5 medium, and then incubated them in a Petri plate. We assessed GFP expression using a fluorescence microscope after 24 or 48 hours.

#### Biolistic delivery

We prepared Biolistic delivery Gold microprojectiles, slightly modifying the method from Hamada *et al*. (2017). To sum up, 5 μg of circular plasmid DNA, 60 mg/mL of Bio-Rad gold particles, 10 μL of 0.1 M spermidine, and 25 μL of 2.5 M CaCl_2_ were mixed together in 50% glycerol. We incubated the mixture for 10 minutes at room temperature, then centrifuged the DNA-coated gold particles at 9,100 g for 2 seconds to pellet them, discarding the supernatant. The pellet was washed sequentially with 70 μL of 70% ethanol, followed by 99.5% ethanol. The particles were washed and then mixed again in 24 μL of 99.5% ethanol. They were sonicated for one second, and then 6 μL liquots were spread out evenly on Bio-Rad macrocarrier membranes and left to dry. The DNA-coated gold particles were delivered using the PDS-1000/He™ particle delivery system at 1350 psi

#### Wheat and barley transformation

We used the binary vectors ICCR1, containing only the BAR gene, and ICCR2, that contains the *BAR* gene and *GRF4-GIF1* chimera, as backbones. The RUBY gene cassette was amplified from the plasmid 160908 (Addgene) and inserted into both the ICCR1 and ICCR2 vectors using the NEBuilder HiFi DNA Assembly Cloning Kit (New England Biolabs, E5520S) to yield the ICCR1:RUBY and ICCR2:RUBY plasmids, respectively. All plasmids were propagated in NEB-stable competent *E. coli* cells (C3040H) and were sequence-verified before being transferred to *A. tumefaciens* strain AGL-1. Transformation of immature embryos of the wheat cultivars Fielder, Chinese Spring, Bobwhite, and Gadish was performed using the ICCR2:RUBY and control ICCR2 constructs, following the method of Shrama *et al*. (2024). Barley transformation was carried out on immature embryos of the cultivar Golden Promise using the ICCR1:RUBY and ICCR2:RUBY constructs, following the procedure of Hinchliffe and Harwood (2019).

#### Determination of transgene DNA copy number

Transgene copy number was determined using ddPCR following previously described protocols (Collier *et al*., 2017; Sharma *et al*., 2023). Genomic DNA was extracted from 4-week-old T0 seedlings using the CTAB method and digested with HaeIII enzyme. The digested DNA served as a template for ddPCR, which was performed using a duplex assay with probe chemistry for the reference gene PUROINDOLINE-b (*PINb*) and the target gene bialaphos resistance (*BAR*) as described in Sharma *et al*. (2024). ddPCR reactions were prepared according to the ddPCR Supermix for Probes kit instructions (Biorad). Droplets were generated using a QX200 droplet generator, and PCR was run on a C1000 deep-well thermal cycler (BioRad). Fluorescence was measured with a QX200 droplet analyzer, and results were evaluated using Bio-Rad Quanta-soft Pro Software.

#### Editing of the CYP76AD1 gene

We designed a sgRNA targeting the *CYP76AD1* gene (Figure 1C). The gRNA was cloned into the JD633 vector (Addgene 160392) using Golden Gate cloning and co-transformed along with the helper vector pRiA4-VIR into *A. tumefaciens* strain *AGL1*. The genome editing plasmid was then transformed into immature embryos of a RUBY-2 transgenic line that contains a single homozygote RUBY cassette. The resulting T0 plants were screened by PCR using the Cas9F and cas9R primers for detection of the *Cas9* gene **(Table S1)**. T1 seeds were harvested from selected green and red pigmented plants, amplified with the CYPF and CYPR *CYP76AD1*-specific primers, and the PCR products were Sanger sequenced. As a control, we amplified and sequenced the *DODA* gene, the second gene in the RUBY cassette.

#### Leaf rust infection assay

Infection with leaf rust was performed according to Khazan et al., (2020). Plants were grown in small pots in a temperature-controlled greenhouse at 22 ±2°C with a 14/10h day night photoperiod. Seedlings were inoculated at one-(7–10 days after planting) or two-leaf stage (10-12 days after planting). Inoculation was performed with urediniospores of *Pt* isolate #526-24. The spores were suspended in light weight mineral oil (Soltrol 170); the oil was allowed to evaporate, and the plants were incubated for 24 h in a dew chamber at 18°C and then moved to a greenhouse and maintained at 22° C for 12–14 days.

#### Salt stress

10-day-old seedlings were irrigated with 70mM NaCl and the concentration was gradually increased to 200 mM by adding 70mM NaCl every second day. The plants were irrigated with 200mM NaCl for additional 14 days and then the phenotype was evaluated by monitoring the shape, size and color of the first (oldest) and last (youngest) emerging leaves.

#### Visualization of fungal infection

Leaves were detached at 8 dpi and stained with WGA-FITC (L4895-10MG; Sigma) as described by Sharma *et al* (2024). The detached leaves were cut into 2-cm pieces and placed in a 10-mL centrifuge tube containing 5 mL of 1 M KOH and 0.05% Silwet L-77. After 12 hours, the KOH solution was gently removed, and the leaves were washed with 10 mL of 50 mM Tris (pH 7.5). This washing step was repeated with another 10 mL of 50 mM Tris (pH 7.5). After 20 minutes, the Tris solution was replaced with 5 mL of 20 μg/mL WGA-FITC. The tissue was stained for 15 minutes and then washed again with 50 mM Tris (pH 7.5). The WGA-FITC-stained tissue was examined under blue light excitation using an Axio Zoom V.16 fluorescent stereoscope (Zeiss) and the area occupied by the fungus was measured.

## Acknowledgements

This work was funded by Israel National Center for Genome Editing in Agriculture

## Author contributions

Plasmid design and construction: D.S., R.K., M.R.P; Protoplast transformation and analysis: R.K., M.R.P.; Generation of transgenic wheat: A.S-C, R.K., M.R.P; Analysis of transgenic lines: A.S-C, M.R.P, D.S, and A.H; CRISPR and analysis of edited lines: M.R.P, A.S-C; Leaf rust phenotyping: M.R.P, A.M.D, and S.E; Salt stress phenotyping: M.R.P.; Conceived study: M.R.P., A.S., A.S-C; Drafted manuscript: M.R.P., A.S.

---

**Figure 1. Genome Manipulation of Wheat Using the RUBY Marker**

Stable transformation of the RUBY gene cassette in wheat cultivar Fielder results in a distinct red phenotype. Subsequent editing of the transgenic RUBY wheat reverts the plants back to their original green phenotype. Along with the loss of the red RUBY phenotype, the edited lines also lose the RUBY-associated traits, such as resistance to fungal infections and tolerance to salt stress.

**Figure 3. Development of RUBY wheat and RUBY barley through betalain overexpression**

(a) Diagram of the transformation plasmids. Top – The RUBY:ICCR2 expression plasmid, bottom – ICCR2 control plasmid with the *BAR* under the *OsAct* promoter (purple), and the *GRF4-GIF1* chimera under the *ZmUbi* promoter (green). The RUBY cassette includes the *CYP76AD1, DODA*, and *GT* genes, separated by the 2A peptides, all under the *Cc35S* promoter. LB and RB denote the left and right borders of the T-DNA region, respectively.

(b) Regeneration of wheat calli transformed with the ICC2:Control(left) or ICCR2:RUBY (right).

(c) RT-PCR analysis of transgenic T0 plants using primers for the *BAR* (995 bp) and *GRF4-GIF1* (1233 bp) gnes; WT - wild type, PC - positive control (ICCR2 plasmid), - NC - negative control (water).

(d) Transgenic wheat lines transformed with the ICC2:Control (left) and the ICCR2:RUBY (right) construct. The RUBY is expressed in all parts of the plants including spikelet and roots.

(e) Barley transformation. Transformation with the ICCR2:RUBY plasmid resulted in red callus but very low regeneration (left), transformation with the ICCR1:RUBY plasmid (without the *GRF4-GIF1* chimera) resulted in high rates of regeneration and transgenic plants.

**Figure 4. Editing of the *CYP76AD1* gene and its effects on betalain production and sensitivity to leaf rust and salt stress.**

(a) Diagram of the gene editing vector JD633, which includes the gRNA sequence targeting the *CYP76AD1* gene. The start site of gRNA is marked in blue, the PAM site in green.

(b) A T0 edited line showing a chimeric phenotype characterized by both red and green coloration.

(c) Whole plant and a spikelet of a RUBY edited T1 plant.

(e) Summary of editing events: “+” denotes insertions, “–” indicates deletion.

(f) Loss of betalain production correlates with reduced resistance in edited lines, as shown by fluorescence images of infection sites at 7 days post-inoculation (dpi), with fungal hyphae stained using WGA- FITC.

(g) Box plot and ANOVA results with Tukey’s pairwise comparison for the pustule colony area. Different letters indicate statistically significant differences (P ≤ 0.01).

(h) Infection of the leaves at 7 dpi. Scale bar in all images =1mm.

(i) Loss of betalain production also leads to decreased salt tolerance in edited lines, as demonstrated by plant phenotypes after 14 days of exposure to 200 mM NaCl.

## Reference

Banakar, R., Schubert, M., Collingwood, M., Vakulskas, C., Eggenberger, A.L. and Wang, K. (2020) Comparison of CRISPR-Cas9/Cas12a Ribonucleoprotein Complexes for Genome Editing Efficiency in the Rice Phytoene Desaturase (OsPDS) Gene. Rice, 13.

Biswal, A.K., Hernandez, L.R.B., Castillo, A.I.R., Debernardi, J.M. and Dhugga, K.S. (2023) An efficient transformation method for genome editing of elite bread wheat cultivars. Front Plant Sci, 14.

Brandt, K.M., Gunn, H., Moretti, N. and Zemetra, R.S. (2020) A Streamlined Protocol for Wheat (Triticum aestivum) Protoplast Isolation and Transformation With CRISPR-Cas Ribonucleoprotein Complexes. Front Plant Sci, 11.

Chen, J., Luo, M., Hands, P., Rolland, V., Zhang, J., Li, Z., Outram, M., Dodds, P. and Ayliffe, M. (2023) A split GAL4 RUBY assay for visual in planta detection of protein–protein interactions. Plant Journal, 114.

Chiu, W.L., Niwa, Y., Zeng, W., Hirano, T., Kobayashi, H. and Sheen, J. (1996) Engineered GFP as a vital reporter in plants. Current Biology, 6.

Collier, R., Dasgupta, K., Xing, Y.P., et al. (2017) Accurate measurement of transgene copy number in crop plants using droplet digital PCR. Plant Journal, 90.

Debernardi, J.M., Tricoli, D.M., Ercoli, M.F., Hayta, S., Ronald, P., Palatnik, J.F. and Dubcovsky, J. (2020) A GRF–GIF chimeric protein improves the regeneration efficiency of transgenic plants. Nat Biotechnol, 38.

Dunbar, T., Tsakirpaloglou, N., Septiningsih, E.M. and Thomson, M.J. (2022) Carbon Nanotube-Mediated Plasmid DNA Delivery in Rice Leaves and Seeds. Int J Mol Sci, 23.

Hamada, H., Linghu, Q., Nagira, Y., Miki, R., Taoka, N. and Imai, R. (2017) An in planta biolistic method for stable wheat transformation. Sci Rep, 7.

Hayta, S., Smedley, M.A., Clarke, M., Forner, M. and Harwood, W.A. (2021) An Efficient Agrobacterium-Mediated Transformation Protocol for Hexaploid and Tetraploid Wheat. Curr Protoc, 1.

He, Y., Zhang, T., Sun, H., Zhan, H. and Zhao, Y. (2020) A reporter for noninvasively monitoring gene expression and plant transformation. Hortic Res, 7.

Jain, G. and Gould, K.S. (2015) Functional significance of betalain biosynthesis in leaves of Disphyma australe under salinity stress. Environ Exp Bot, 109.

Jin, S. xia, Liu, G.ze, Zhu, H.guo, Yang, X.yan and Zhang, X.long (2012) Transformation of Upland Cotton (Gossypium hirsutum L.) with gfp Gene as a Visual Marker. J Integr Agric, 11.

Jung, M., Shin, S.H., Park, J.M., Lee, S.N., Lee, M.Y., Ryu, K.H., Paek, K.Y. and Harn, C.H. (2011) Detection of transgene in early developmental stage by GFP monitoring enhances the efficiency of genetic transformation of pepper. Plant Biotechnol Rep, 5.

Khazan, S., Minz-Dub, A., Sela, H., Manisterski, J., Ben-Yehuda, P., Sharon, A. and Millet, E. (2020) Reducing the size of an alien segment carrying leaf rust and stripe rust resistance in wheat. BMC Plant Biol, 20.

Liang, Z., Chen, K., Zhang, Y., Liu, J., Yin, K., Qiu, J.L. and Gao, C. (2018) Genome editing of bread wheat using biolistic delivery of CRISPR/Cas9 in vitro transcripts or ribonucleoproteins. Nat Protoc, 13.

Lin, C.S., Hsu, C.T., Yang, L.H., et al. (2018) Application of protoplast technology to CRISPR/Cas9 mutagenesis: from single-cell mutation detection to mutant plant regeneration. Plant Biotechnol J, 16.

Miki, B. and McHugh, S. (2004) Selectable marker genes in transgenic plants: Applications, alternatives and biosafety. J Biotechnol, 107.

Najafi, S., Bertini, E., D’Incà, E., Fasoli, M. and Zenoni, S. (2023) DNA-free genome editing in grapevine using CRISPR/Cas9 ribonucleoprotein complexes followed by protoplast regeneration. Hortic Res, 10.

Omar, A.A., Song, W.-Y., Graham, J.H. and Grosser, J.W. (2019) Comparison between Protoplast Transformation and Co-transformation in ‘Hamlin’ Sweet Orange [Citrus sinensis (L.) Osbeck]. HortScience, 41.

Pierroz, G. (2023) Seeing red: a modified RUBY reporter assay for visualizing in planta protein–protein interactions. Plant Journal, 114.

Polturak, G., Grossman, N., Vela-Corcia, D., Dong, Y., Nudel, A., Pliner, M., Levy, M., Rogachev, I. and Aharoni, A. (2017) Engineered gray mold resistance, antioxidant capacity, and pigmentation in betalain-producing crops and ornamentals. Proc Natl Acad Sci U S A, 114.

Rather, G.A., Ayzenshtat, D., Teper-Bamnolker, P., Kumar, M., Forotan, Z., Eshel, D. and Bocobza, S. (2022) Advances in protoplast transfection promote efficient CRISPR/Cas9-mediated genome editing in tetraploid potato. Planta, 256.

Sharma D A,.., Avni, R., Gutierrez-Gonzale, J et al. (2023) A single NLR gene confers resistance to leaf and stripe rust in wheat. Res Sq.

Siddappa, S., Sharma, N., Salaria, N., et al. (2023) CRISPR/Cas9-mediated editing of phytoene desaturase (PDS) gene in an important staple crop, potato. 3 Biotech, 13.

Sparkes, I.A., Runions, J., Kearns, A. and Hawes, C. (2006) Rapid, transient expression of fluorescent fusion proteins in tobacco plants and generation of stably transformed plants. Nat Protoc, 1.

Tanaka, J., Minkenberg, B., Poddar, S., Staskawicz, B. and Cho, M.J. (2022) Improvement of Gene Delivery and Mutation Efficiency in the CRISPR-Cas9 Wheat (Triticum aestivum L.) Genomics System via Biolistics. Genes (Basel), 13.

Tassy, C., Partier, A., Beckert, M., Feuillet, C. and Barret, P. (2014) Biolistic transformation of wheat: increased production of plants with simple insertions and heritable transgene expression. Plant Cell Tissue Organ Cult, 119.

Wang, D., Zhong, Y., Feng, B., et al. (2023) The RUBY reporter enables efficient haploid identification in maize and tomato. Plant Biotechnol J, 21.

Yang, S.H., Kim, S.W., Lee, S. and Koo, Y. (2024) Optimized protocols for protoplast isolation, transfection, and regeneration in the Solanum genus for the CRISPR/Cas-mediated transgene-free genome editing. Appl Biol Chem, 67.

Yu, Y., Yu, H., Peng, J., et al. (2024) Enhancing wheat regeneration and genetic transformation through overexpression of TaLAX1. Plant Commun, 5.

Zhang, Z., Hua, L., Gupta, A., Tricoli, D., Edwards, K.J., Yang, B. and Li, W. (2019) Development of an Agrobacterium-delivered CRISPR/Cas9 system for wheat genome editing. Plant Biotechnol J, 17.

